# Dual avatars of *E. coli grxB* encoded Glutaredoxin 2 perform ascorbate recycling and ion channel activities

**DOI:** 10.1101/2021.08.28.458008

**Authors:** Sreeshma Nellootil Sreekumar, Bhaba Krishna Das, Rahul Raina, Neethu Puthumadathil, Sonakshi Udinia, Amit Kumar, Sibasis Sahoo, Pooja Ravichandran, Suman Kumar, Pratima Ray, Dhiraj Kumar, Anmol Chandele, Kozhinjampara R. Mahendran, Arulandu Arockiasamy

## Abstract

Glutaredoxins (Grxs) are single-domain redox enzymes of the thioredoxin superfamily, and primarily function as glutathione (GSH) dependent disulphide reductases. Whereas, the *E. coli* Glutaredoxin 2 (*Ec*Grx2) encoded by *grxB* has two conserved GST-fold domains, it still lacks a classical Grx-like functions. In this study, we show for the first time, that *Ec*Grx2 exists in both soluble and membrane integrated forms. The soluble form associates with a previously unidentified GSH dependent dehydroascrobate (DHA) reductase, and the membrane integrated form possesses ion channel activities. Using enzyme kinetic data and structural data we unequivocally demonstrate that *Ec*Grx2 recycles ascorbate (AsA) from DHA. This ability to recycle AsA is inhibited by Zinc (Zn^2+^). We also show that both wildtype and the *E. coli grxB* deletion mutant can be rescued from H_2_O_2_-induced oxidative stress using ascorbate as an antioxidant, which otherwise is only known as a carbon source in bacteria. Moreover, the *grxB*^*-*^ mutant is susceptible to intracellular killing by ROS producing macrophages. We further discovered that *Ec*Grx2 integrates into the native *E. coli* membrane and show that the purified soluble protein readily inserts into artificial lipid bilayer membrane and conducts ions *in vitro*. Our data demonstrates a highly conserved functional similarity among *Ec*Grx2-orthologs and highlights that the utilization and subsequent recycling of ascorbate as an antioxidant by *grxB* harbouring gram-negative bacteria, including human pathogens, may provide a survival advantage under hostile oxidative environments.

## Introduction

Omnipresent Grxs are involved in various biological functions such as reduction of ribonucleotide reductase (RNR), phosphoadenosine phosphosulfate (PAPS) reductase, and arsenate reductase, cell cycle control, signal transduction, and dehydroascorbate reductase activities^1-4^. However, *Ec*Grx2 is atypical and has a two-domain architecture^5,6^ compared to the single-domain Grx1, 3 and 4 of *E. coli*^7,8^. The biochemical role of this duplication is unclear. While *Ec*Grx2 lacks key RNR reduction activity, it reduces protein and mixed disulphides in ArsC^4^ and PAPS reductase. Notably, *Ec*Grx2 is abundantly expressed during stationary phase, upregulated under osmotic shock, and acidic conditions^9,10^. Thus, predictably, null mutants (*grxB*^-^) were reported to show distorted morphology and an increased susceptibility under H_2_O_2_-induced oxidative conditions^4,11^. Cumulatively, these reports strongly suggest *Ec*Grx2 as an antioxidant enzyme. Here, we employ a structure-based approach to decipher the precise biochemical functions associated with the three-dimensional structure of *Ec*Grx2.

## Results

### Soluble *Ec*Grx2 is a GSH-dependent DHA reductase

To comprehensively elucidate its biochemical activities, we first compared the crystal structure of *Ec*Grx2 with that of its eukaryotic structural homologs. The structure of monomeric *Ec*Grx2^5,7^ has a N-terminal domain (NTD; GST_N_3) containing βαβαββα thioredoxin-fold^12^, and an all α-helical C-terminal domain (CTD; Glutaredoxin2_C)^13^ (Fig. 1a). Based on the structural similarity search, we retrieved the *Pennisetum glaucum* dehydroascrobate reductase (*Pg*DHAR)^14^ and human chloride intracellular channel CLIC1 (*Hs*CLIC1) with high Z-score of >14.4 in DALI searches, in spite of the low sequence identity (<20%). The overall structure of *Ec*Grx2, *Pg*DHAR, and *Hs*CLIC1 are highly conserved (RMSD ≤3.6 Å). It is noteworthy that both plant DHARs and *Hs*CLICs are redox enzymes that recycle AsA from DHA using GSH as a cofactor^15,16^. Plant DHAR is part of the well-studied AsA-GSH based antioxidant system (Foyer-Halliwell-Asada pathway) that helps recover from environmental stress^17^. The enzymatic activity of *Pg*DHAR and *Hs*CLIC1 is imparted by a catalytic cysteine present in the conserved glutaredoxin motif: Cxx[C/S] (Fig. 1b). Further, the active site in these structural orthologs is composed of canonical GSH binding ‘G-site’ at the NTD, and a hydrophobic substrate binding ‘H-site’ from the CTD^14^. A closer inspection of the structure-guided multiple sequence alignment and the electrostatic surface analysis reveals a distinct positively charged pocket conserved in *Ec*Grx2 with a CxxC motif (Cys9-Pro10-Tyr11-Cys12) (Fig. 1c). These structural and active site similarities lead us to hypothesize that soluble *Ec*Grx2 may possess the ability to recycle AsA from DHA like its eukaryotic structural orthologs.

**Fig. 1.**
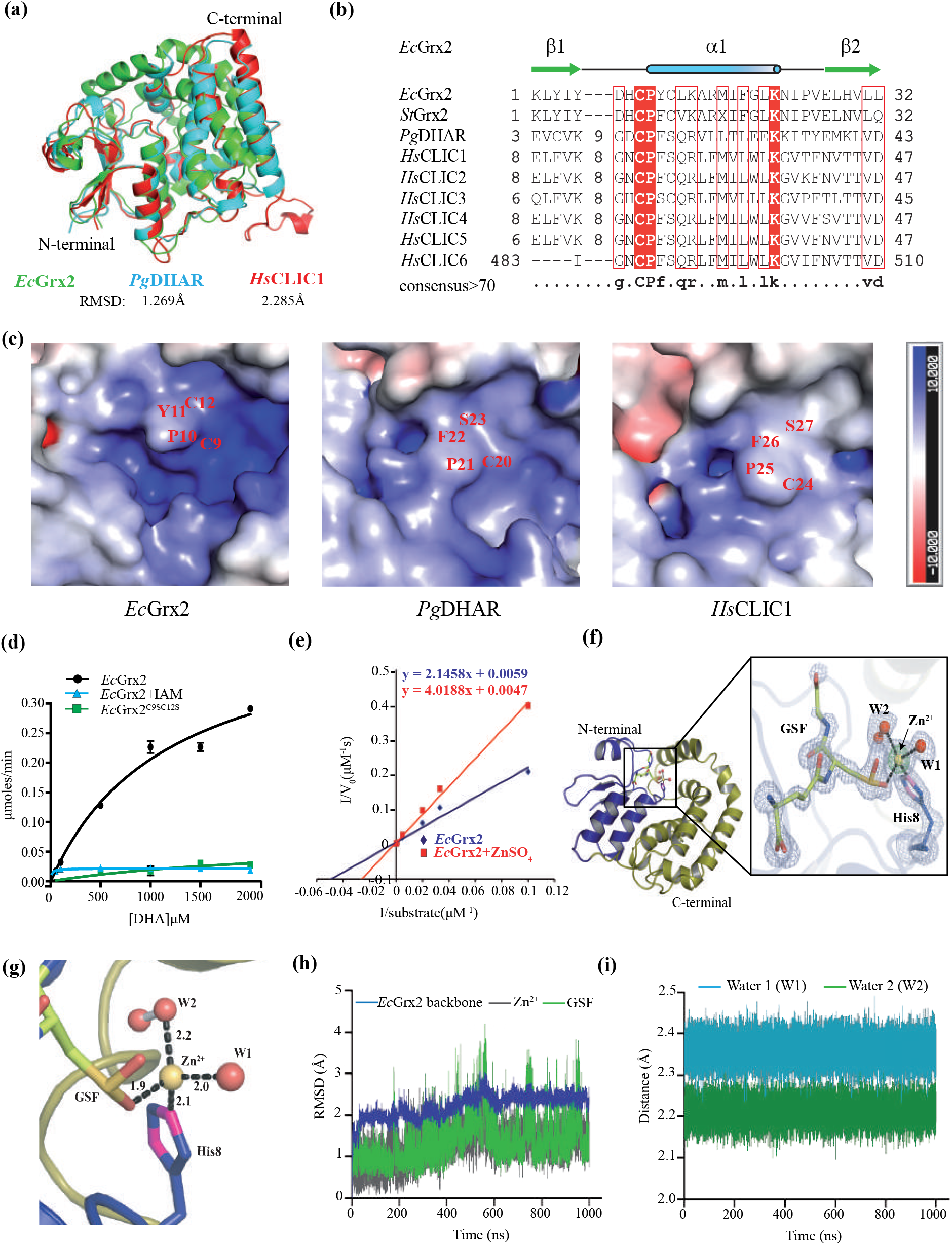
*Ec*Grx2 is a DHA reductase and Zn^2+^ ion inhibits enzyme activity. **(a)** Structural superposition of *Ec*Grx2 (7DKP, green) and *Pg*DHAR (5EVO, cyan) and *Hs*CLIC1 (1K0M, red). **(b)** Structure-based multiple sequence alignment showing conserved active site motif: CxxC(S). *Pg: Pennisetum glaucum, Hs: Homo sapiens, Ec: Escherichia coli, St: Salmonella typhimurium*, **(c)** Electrostatic potential surface maps of predicted active site region in *Ec*Grx2 in comparison to *Pg*DHAR and *Hs*CLIC1 showing conserved residues. **(d)** Michaelis-Menten kinetics* of *Ec*Grx2; native, double mutant and chemically modified protein is shown. *Ec*Grx2(●), *Ec*Grx2-reduced and treated with IAM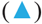, *Ec*Grx2^C9S/C12S^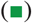. **(e)** Lineweaver-Burk plot* showing competitive inhibition by Zn^2+^. *Ec*Grx2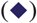, *Ec*Grx2 + 45 μM ZnSO_4_ 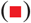, **(f)** Crystal structure of Zn^2+^ inhibited *Ec*Grx2 with the inset showing omit difference (Fo-Fc) map contoured at 3σ (grey), wherein Zn^2+^ ion is shown at 15σ (green). **(g)** Tetrahedral coordination of Zn^2+^ with GSF, two waters (W1, W2) and His8 is shown. **(h)** RMSD plot of 1 µs MD simulation of *Ec*Grx2-GSF-Zn^2+^ complex is shown for protein backbone (Blue), GSF (Green) and Zn^2+^ (Grey). **(i)** Distance plot representing interatomic distance between Zn^2+^ and the two coordinating waters are shown. *Kinetic values represent mean ± standard deviation of technical triplicates (n= 3). AsA monitored (min^-1^) at A _265_ is plotted against different concentration of DHA (μM).

Therefore, we then asked whether *Ec*Grx2 recycles AsA from DHA, like *Pg*DHAR and *Hs*CLIC1^16,17^. As we hypothesized, the purified *Ec*Grx2 showed GSH-dependent DHA reductase activity at an optimal pH of 7.4, following Michaelis-Menten kinetics (Supplementary Fig. 1 a, b, c and Supplementary Table 1). The K_m(DHA)_, K_cat(DHA)_ and specific activity were 2879 ± 640.4 µM, 229.47 min^-1^ and 1.8845 μM^-1^min^-1^mg^-1^, respectively. We then probed to find the role of Cys9 and Cys12 in DHA reductase activity after chemical modification by carbamidomethylation^18^, which abrogated the enzyme activity completely (Fig. 1d). We then substantiated this observation by creating a double mutant (C9S/C12S) that resulted in loss of enzyme activity (Fig. 1d). The above data clearly shows that *Ec*Grx2 recycles AsA from DHA in a GSH-dependent manner. As thiol-reactive divalent cations are known to inhibit DHA reductase activity of plant DHAR^15^, we performed a thermal shift assay (TSA) to screen for divalent cations that inhibit *Ec*Grx2 enzyme activity. In contrast to other divalent metal ions that did not show significant effect on melting temperature (T_m_), except Cu^2+^ (Supplementary Fig. 1d), addition of Zn^2+^ ion resulted in a decrease in T_m_ by 7 °C, implying direct binding to *Ec*Grx2. Further, we also observed that Zn^2+^ abrogates DHA reductase activity of *Ec*Grx2, competitively (Fig. 1e). Then, in order to understand the molecular mechanism of how Zn^2+^ inhibits DHA reductase activity, we determined the first crystal structure of *Ec*Grx2 in complex with Zn^2+^ and GSH at 1.6 Å (Table 1, PDB:7D9L). The complex structure revealed a hitherto unknown mode of Zn^2+^ inhibition at the enzyme active site, wherein a highly conserved histidine (His8), two water molecules, and the cysteine of oxidised GSH (GSOO:GSF) form a tetrahedral coordination^19^ with Zn^2+^ (Fig. 1f,g and Supplementary Table 2). Interestingly, Zn^2+^ is coordinated by GSF, instead of canonical thiol (-SH) group of catalytic cysteine. More so, the bound GSH is oxidised to sulfinic acid form (GSOO) by the Zn^2+^ at the G-site (Supplementary Fig. 2a). Moreover, there is no significant change noticed in the overall structure of *Ec*Grx2 among native (2.38 Å), GSH bound (1.45 Å) and Zn^2+^ inhibited crystal structures reported here (Table 1, PDB: 7DKR, 7DKP), and by others (PDB:4KX4, 3IR4). Further, we performed 1 µs molecular dynamics simulation at 300K and found that the Zn^2+^ inhibited complex is very stable with an RMSD of GSF;1.36 ± 0.52 Å, Zn^2+^; 1.31 ± 0.42, water-1; (2.2 ± 0.03 Å) and water-2; 2.21 ± 0.02 Å (Fig. 1 h, i, Supplementary Fig. 2 b, Supplementary video 1). In addition to the active site bound Zn^2+^, a second Zn^2+^ ion is present in the crystal structure which is tetrahedrally coordinated by symmetry-related *Ec*Grx2 monomer involved in crystal packing (Supplementary Fig. 2c, d). The above data unequivocally demonstrate that the AsA recycling DHA reductase activity of *Ec*Grx2 involves the conserved CxxC motif and the divalent cation Zn^2+^ inhibits the enzyme activity by binding at the G-site, prompting us to probe whether AsA is used as an antioxidant by *E. coli*.

**Table 1.** X-ray data collection and refinement statistics.

| **Data collection (PDB)** | ***Ec*Grx2-apo (7DKR)** | ***Ec*Grx2-GSH (7DKP)** | ***Ec*Grx2-Zn<sup>2+</sup>-GSF (7D9L)** |
| --- | --- | --- | --- |
| X ray wavelength (Å) | 1.00 | 0.96770 | 0.98400 |
| Space group | P 2<sub>1</sub>2<sub>1</sub>2<sub>1</sub> | P 1 2<sub>1</sub> 1 | P 2<sub>1</sub>2<sub>1</sub>2 |
| **Cell dimensions** |  |  |  |
| a, b, c (Å) | 28.47, 78.95, 89.28 | 49.827, 169.408, 49.836 | 46.095, 106.758, 42.130 |
| $\alpha$, $\beta$, $\gamma$ (°) | 90.0, 90.0, 90.0 | 90.0, 93.52, 90.0 | 90.0, 90.0, 90.0 |
| Resolution (Å) | 44.64 - 2.38 (2.47 - 2.38) | 42.893 - 1.449 (1.476 - 1.449) | 42.319 - 1.60 (1.628 - 1.60) |
| Refs. total/unique | 95444/8647 | 576328/137631 | 176283/28045 |
| R<sub>merge</sub> (%) | 0.111 (0.297) | 0.099 (0.635) | 0.19 (1.207) |
| I/$\sigma$ (I) | 15.2 (6.5) | 8.7 (1.9) | 11.7 (3.4) |
| Completeness (%) | 100 (100) | 94.7 (89.0) | 99.3 (99.4) |
| Redundancy | 11.0 (9.5) | 4.2 (4.2) | 6.3 (6.2) |
| CC<sub>1/2</sub> | 0.997 (0.973) | 0.997 (0.759) | 0.995 (0.555) |
| **Refinement** |  |  |  |
| Resolution (Å) | 2.38 | 1.45 | 1.61 |
| R<sub>work</sub>/R<sub>free</sub> (%) | 0.195/0.249 | 0.138/0.160 | 0.130/0.182 |
| **No. of atoms** |  |  |  |
| Protein (non-H atoms) | 1682 | 6906 | 1694 |
| L-gamma-glutamyl-3-sulfino-L-alanyl-glycine | - | - | 22 |
| γ-L-Glutamyl-L-cysteinylglycine | - | 80 | - |
| Oxygen molecule | - | - | 2 |
| Zinc ion | - | - | 2 |
| Chloride ion | - | - | 1 |
| Sulfate ion | - | - | 5 |
| Citrate anion | - | 39 | - |
| Water | 104 | 1314 | 156 |
| **Average B factor (Å<sup>2</sup>)** |  |  |  |
| Protein | 20.839 |  | 14.332 |
| Water | 24.487 | 25.962 | 27.212 |
| **r.m.s. deviations** |  |  |  |
| Bond lengths (Å) | 0.43 | 0.91 | 0.82 |
| Bond angles (°) | 0.55 | 1.02 | 1.06 |
| **Ramachandran plot** |  |  |  |
| Favoured | 205 (96%) | 847 (98%) | 211 (98%) |
| Allowed | 7 (3%) | 19 (2%) | 5 (2%) |
| Outliers | 1 (0%) | 3 (0%) | 0 |

r.m.s., root mean square. Each data set was collected from a single crystal. Values in parentheses are for high-resolution shell.

### Ascorbate rescues *E. coli* from H_2_O_2_ induced oxidative stress

AsA has been previously studied as a carbon source in bacteria^20-22^ and there is no evidence for its use as an antioxidant by *E. coli*. Therefore, we performed AsA supplementation assay with wild type *E. coli* (BW25113) and *Ec*Grx2 deletion mutant (JW1051-3: *ΔgrxB734::kan*) strains, using M9 minimal media-agar plates containing varying concentrations of AsA (0.5-15.1 mM) under H_2_O_2_ induced oxidative stress (Fig. 2a, b, Supplementary Table 3). When AsA was not supplemented, the zone of clearance was larger in Grx2 mutant compared to that of wild type strain, suggesting that the *grxB* deletion mutation increases bacterial sensitivity to H_2_O_2_, in concurrence with the earlier observation^4^. We further observed a concentration dependent decrease in zone of clearance upon AsA supplementation in both wild type and mutant strains, indicative of an antioxidant role played by AsA in minimal media devoid of redox active chemicals.

**Fig. 2.**
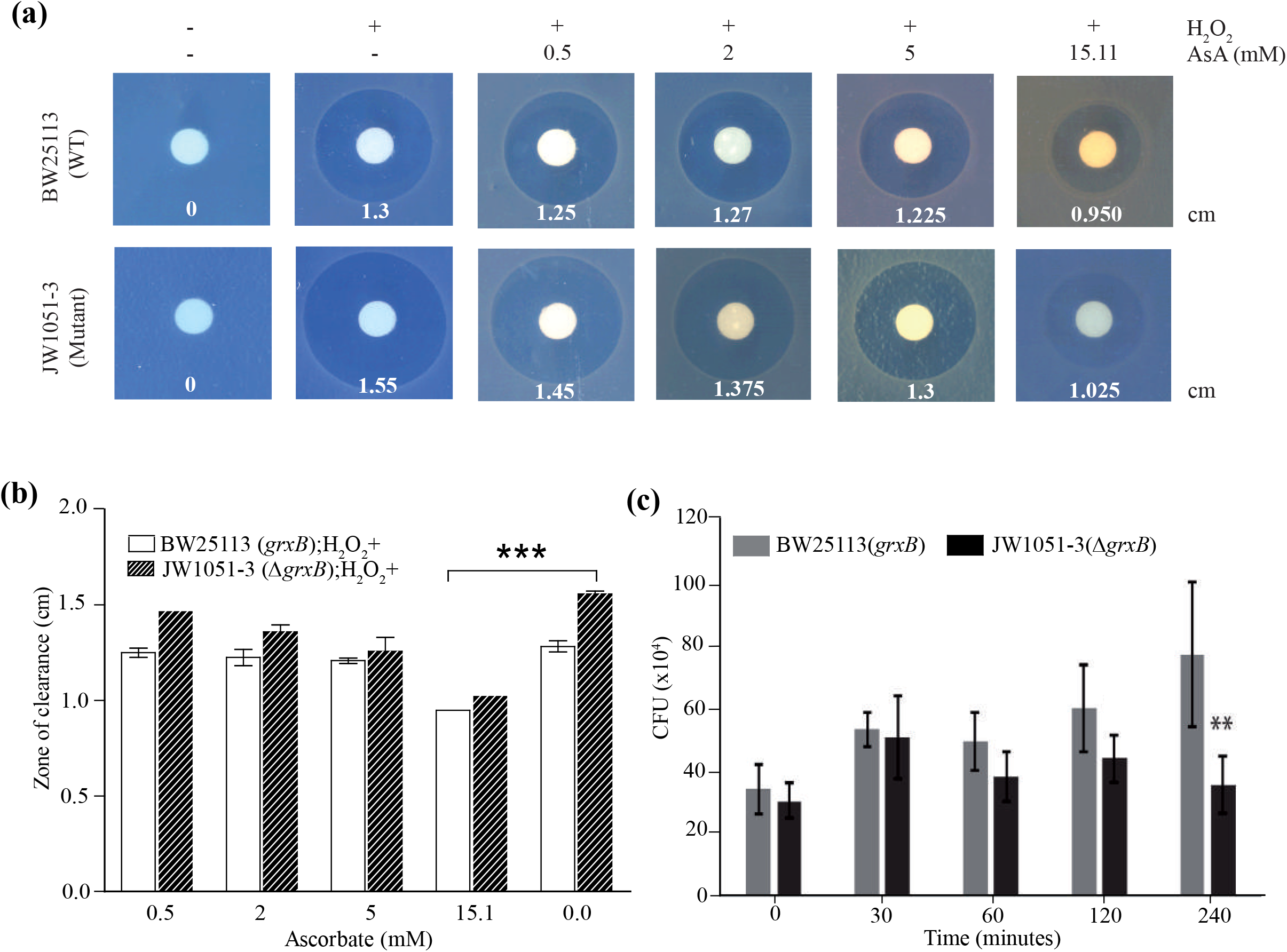
AsA rescues *E. coli* from H_2_O_2_ induced stress and *Ec*Grx2 mutant is susceptible to macrophage mediated killing. **(a, b)** ascorbate supplementation assay using filter disc saturated with H_2_O_2_ is shown using wild type and mutant strains BW25113, JW1051-3 (*ΔgrxB*, Kan^R^), respectively. Zone of clearance (cm) was calculated from triplicates (biological replicates, n=2), and statistical significance calculated using one-way ANOVA (*, p < 0.05). **(c)** Survival of wild type and mutant *E. coli* strains inside LPS activated RAW264.7 murine macrophages is represented as number of CFU/ml, plotted at different time points. Bars = mean ± s.e.m., values are from three independent experiments. *Significantly different from WT (p <0.001) using one-way ANOVA.

### *Ec*Grx2 provides bacteria survival advantage inside LPS activated RAW264.7 macrophages

As the *grxB*^-^ mutant shows phenotypic changes only in the presence of oxidising environment^9^, we asked whether *Ec*Grx2 helps bacteria survive inside the immune cells that produce more ROS upon bacterial infection, as predicted before^9^. We then screened both wild type and *grxB* mutant strains for their ability to survive inside the host cell (RAW264.7 macrophages) upon infection. The *grxB* mutant *E. coli* showed a significant decrease in survival (*P* <0.001) compared to the wild type strain (Fig. 2c, Supplementary Table 4), indicating a possible antioxidant role for *Ec*Grx2 inside the host macrophages. Having now demonstrated DHA reductase activity for the soluble form of *Ec*Grx2, we further asked whether the protein is also integrated into the membrane, similar to its eukaryotic orthologs *Hs*CLICs^23-25^ and plant DHAR^26^.

### *Ec*Grx2 moonlights as an ion channel

To understand the dimorphic nature of *Ec*Grx2, we probed the dual localization of *Ec*Grx2-eGFP in wild type *E. coli* (BW25113) under stationary phase conditions during which maximal expression was expected. Confocal microscopy followed by TIRF-SIM analysis clearly show that *Ec*Grx2-eGFP is localized into both cytoplasmic and membrane compartments whereas eGFP alone is confined to the cytoplasm (Fig. 3a (i-v)). Membrane integration of *Ec*Grx2-eGFP was further assessed by isolating pure membrane fraction from *E. coli*, and subsequent extraction with various detergents with SDS as control. The membrane fraction was first washed with 2 M urea, before detergent extraction, to exclude the peripherally attached soluble *Ec*Grx2 from the membrane integrated form. We show that *Ec*Grx2 is present in the urea washed membrane and also in the detergent extracted supernatant (Fig. 3b), confirming that *Ec*Grx2 is integrated into the native *E. coli* membrane. Then, we sought to determine if the soluble form of *Ec*Grx2 inserts spontaneously into the planar lipid bilayer and forms a functional ion channel similar to that of *Hs*CLICs^27,28^.

**Fig. 3.**
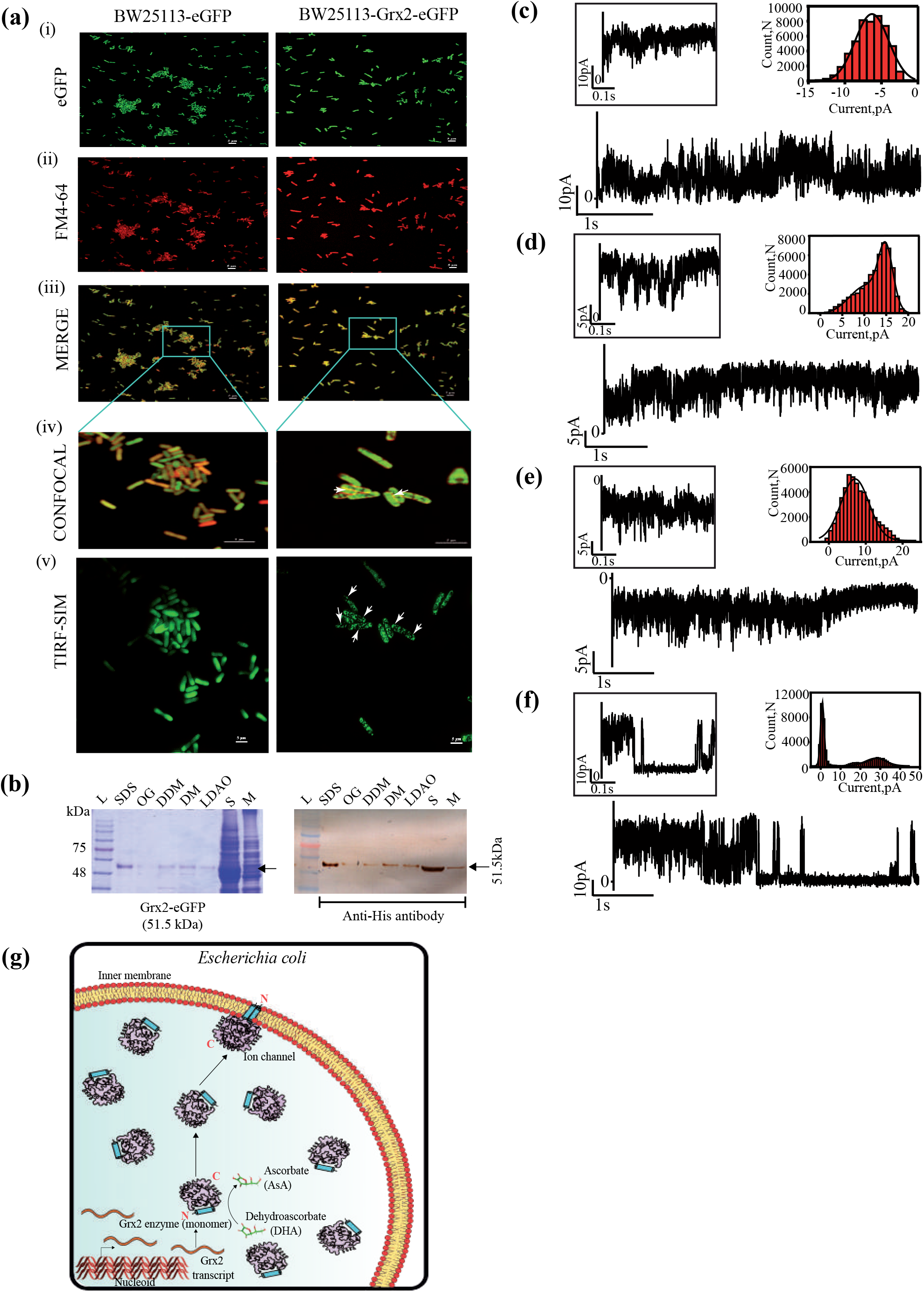
*Ec*Grx2 exists in membrane integrated form and soluble protein conducts ion after inserting into lipid bilayer. **(a)** Confocal and TIRF-SIM microscopic images showing membrane localization of Grx2 in BW25113 E. coli cells. left: cells transformed with eGFP alone, right: cells transformed with Grx2-eGFP. (i) eGFP channel, (ii) stained for plasma membrane, (iii) merged images of (i) and (ii), and the inset (iv) showing 4 X zoom and the corresponding region under TIRF-SIM (v) is shown with arrows pointing to the membrane localized protein. Scale bars=5µM. **(b)** 4-20% reduced SDS-PAGE (left) and corresponding Western blot (right) loaded with detergent extracted samples using pure membranes isolated from BL21 (DE3). L: standard markers, S: ultra-supernatant, M: membrane after 2M urea wash. Uncropped gel and blot are given in supplemental information. **(c-f)** Ion current recordings of *Ec*Grx2 reconstituted into planar lipid bilayers at **(c)** +50 mV, **(d)** flickering events characteristic of channels at +50 mV, **(e)** -50 mV showing noisy current fluctuations, **(f)** channel gating at +100 mV. Buffer conditions: 1 M KCl, 10 mM MES, pH 6.0. The current signals were filtered at 2 kHz and sampled at 10 kHz. **(g)** Model depicting the dimorphic and bifunctional nature of *Ec*Grx2.

The channel-forming properties such as ion conductance and gating of *Ec*Grx2 were examined using single-channel electrical recordings. Planar lipid bilayer were made by the Montal and Muller technique^29^ using DPhPC (1, 2-diphytanoyl-sn-glycero-3-phosphocholine) lipids that provide highly stable membranes. Soluble *Ec*Grx2 inserts into the lipid bilayers at + 50 mV, forming channels that fluctuated between different conductance states, indicating the formation of non-uniform channels in the membrane (Fig. 3c). Notably, the channel formation is associated with noisy transient spikes of varying conductance states and the channel remained in an open conductance state for a longer duration at + 50 mV (Fig. 3c, d). Importantly, channel activity is evident from the short ion current flickering events corresponding to the open and closed conductance states. At - 50 mV, the channel showed numerous transient openings and fluctuating conductance states with increased noise leading to rapid closure of the channel (Fig. 3e). Further, we observed asymmetry in the channel conductance states and fluctuation with respect to the channel with the direction of the applied voltage. At higher voltages of +100 mV, *Ec*Grx2 exhibited voltage-dependent gating subsequently resulting in the complete channel closure (Fig. 3f). The frequency of gating events increased with the increase in the voltage and the time duration of open and closed states varied from several seconds to minutes. Our single-channel electrical recordings confirm the channel formation of *Ec*Grx2, which exhibited consistent electrical properties such as ion conductance and gating in a membrane environment. The above data demonstrate that *Ec*Grx2 is membrane integrated and the soluble form readily inserts into artificial bilayer membrane and conducts ions.

Taken together, our data show that a) the atypical two domain containing *Ec*Grx2 is a GSH dependant DHA reductase that recycles AsA from DHA in its soluble monomeric form; b) Zn^2+^ ion inhibits *Ec*Grx2 enzyme activity; c) The complex crystal structure followed by molecular dynamics simulation analysis shows Zn^2+^ ion forms a stable inhibitory complex through tetrahedral coordination geometry at the G-site; d) AsA, as an antioxidant, rescues *E. coli* cells from H_2_O_2_ induced stress *in vitro*; e) *E. coli grxB*^*-*^ mutant is susceptible to increased killing by RAW macrophages upon infection; f) *Ec*Grx2 is dimorphic and has a moonlighting membrane integrated form (Fig. 3g); and g) The soluble form of *Ec*Grx2 inserts into artificial bilayer membrane and forms transient ion channels, similar to *Hs*CLICs.

## Discussion

Eukaryotes use AsA as a major antioxidant and thus recycling from either monodehydroascorbate (MDHA) radical and/or DHA is essential for cellular survival and alleviation of oxidative stress induced damage. Though AsA utilization and DHA reduction by *E. coli* has been studied for some time^20,30,31^, the antioxidant role of AsA and its enzyme-mediated recycling remained unexplored till now. The identity of DHA reductase activity reported for partially purified fraction of *E. coli* extract^32^ remains unknown. Thus, GSH dependent AsA recycling activity reported here shows that *Ec*Grx2 may well be a part of the hitherto unknown bacterial antioxidant defence system. Interestingly *grxB* gene is widely distributed among Gram-negative proteobacteria (Supplementary Fig. 3 a, b). The extensive parallels; dimorphic nature (Fig. 3g), sub-cellular localization^33,34^, and dual biochemical activities conserved among *Ec*Grx2, *Hs*CLICs and plant DHARs support the above hypothesis. Further studies are needed to assess whether the moonlighting membrane integrated ion channel form of *Ec*Grx2 also plays a protective role under oxidative stress. As *grxB* is required for late stages of bacterial growth and survival inside macrophages, *Ec*Grx2-orthologs from pathogenic bacteria are potential drug targets for therapeutics that could function by disrupting the microbial antioxidant defense system.

## Materials and Methods

### Materials

Unless otherwise indicated all chemicals and reagents were purchased from Sigma–Aldrich (Merck). Restriction enzymes were from New England Biolabs. Oligonucleotides were from Macrogen. Bacterial culture media were from Himedia. Strains BW25113 (CGSC#:7636) and JW1051-3 (CGSC#9012)^35^ were obtained from *E. coli* Genetic Stock Center, Yale university. Cell culture materials and media are from Nunclon, Cell clone, Gibco and Merck.

### Bacterial strains, plasmids, primers and culture conditions

Details are summarized in Supplementary Table 5. *E. coli* strains were grown with shaking at 130 rpm in Luria-Bertani (LB) or in LB auto induction (LB-AIM)^36^ media without trace elements (Formedium) or in 1.5% LB agar plates. The following concentrations of antibiotics were used: Ampicillin 100 µgml^-1^, Chloramphenicol 34 µgml^-1^, Kanamycin 50 µgml^-1^. *E. coli* DH5a was used as the host strain for all plasmid constructions.

### Plasmid construction

pETM11-*grxB*, encoding His6-*Ec*Grx2, was generated by PCR amplification of *E. coli grxB* from pET24a-*grxB* (gifted by H. Jane Dyson, The Scripps Research Institute) and cloned into pETM11 (EMBL). pETM11-*grxB* ^C9S/C12S^ and pETM1-*grxB*^C9S^, encoding His6-*Ec*Grx2^C9S/C12*S*^ and His6-*Ec*Grx2^C9S^; double and single mutants, were made by PCR amplification using mutagenic primers and pETM11-*grxB* as template. pQE30-*grxB* encoding His6-*Ec*Grx2 was generated using pETM1-*grxB* as template for PCR amplification. pET16b-*grxB*-*eGFP*, encoding *Ec*Grx2-eGFP-His6, was generated by amplifying *grxB* from pET24a-*grxB* and cloned into pET16b-eGFP (gifted by Aravind Penmatsa, MBU, IISc, Bangalore). pQE30-*grxB-eGFP*, encoding His6-*Ec*Grx2-eGFP, was generated by PCR amplification of *grxB-eGFP* from pET16b-*grxB-eGFP*. Plasmid pQE30-*eGFP*, encoding His6-eGFP, was generated by PCR amplification of *eGFP* from pET16b-*eGFP*.

### Protein expression and purification

*EcGrx2(native):* pET24a encoding tagless native *Ec*Grx2 was transformed into *E. coli* Rosetta2 (DE3) pLysS and overexpression was optimized in LB-AIM. Briefly, cells from single colony were inoculated for primary culture in 25 ml LB broth. Secondary culture was raised from 1% of the primary inoculum at 37 °C at 130 rpm to an OD_600_ of 0.6, supplemented with Kanamycin and Chloramphenicol. The culture was grown at 25 °C, 130 rpm for 24 hours. Cells were harvested and resuspended in ice-cold buffer-A (Supplementary Table 6) and were disrupted by sonication at room temperature (RT). Cell lysate was centrifuged at 15000 g for 30 minutes at 4 °C and the supernatant fraction was 0.45 μm filtered. The filtrate was dialyzed against 10 mM sodium acetate buffer pH 4.5-5.0 containing 10mM NaCl and centrifuged at 18900 x g for 10 minutes at 4 °C. Supernatant obtained after centrifugation was directly loaded onto 5 ml HiTrap-SP FF cation-exchange column (GE) equilibrated with buffer-B. Protein was eluted over a linear gradient of 10-100mM NaCl. Cleaner fractions were pooled and concentrated using ultracentrifugation device; Centriprep (3kDa, Amicon). Protein was passed through Superdex S75 16/60 column (GE) equilibrated with buffer-C. Peak fractions were pooled and dialyzed against buffer-D. Peak fractions were pooled and concentrated to 35 mg/ml. Protein purity was assessed using a 4-20% SDS-PAGE and concentration were estimated using OD280nm measurement and theoretical molar extinction coefficient (22920 M^-1^cm^-1^). Prior to storage, dithiothreitol (DTT) was added to final concentration of 10 mM. 50 μl aliquots in PCR vials were flash frozen in liquid nitrogen (LN2) and stored at -80 °C till further use. *EcGrx2*^*C9S/C12S*^ *and EcGrx2*^*C9S*^: The following protocol was followed for overexpression and purification of both *Ec*Grx2^C9S/C12S^ and *Ec*Grx2^C9S^ mutants. Briefly, plasmids pETM11-*grxB*^C9S/C12S^ and pETM11-*grxB*^C9S^ were transformed into *E. coli* BL21(DE3).

Overexpression and cell lysis were carried out similar to native *Ec*Grx2. The 0.45 µm filtered supernatant was passed through 5ml HisTrap FF column (GE) equilibrated with buffer-E. The column was washed with 10 cv of buffer-E followed by elution using a linear gradient of 10-300 mM Imidazole. Peak fractions were pooled and concentrated using centriprep and was passed through Superdex S75 16/60 column (GE) equilibrated with buffer-F. Peak fractions containing pure protein were pooled, concentrated to 4 mg/ml and 50 μl of aliquots were flash frozen in LN2 and stored at -80 °C. *EcGrx2-eGFP-His6*: Plasmid pQE30-*grxB-eGFP* was transformed into DH5α and tested for protein expression prior to microscopic studies. Briefly, cell lysis was carried out similar to native *Ec*Grx2 and the lysate was centrifuged at 15000 g for 30 minutes at 4 °C and then the supernatant was used for binding Ni-NTA beads, eluted with buffer-G and checked on SDS-PAGE. *eGFP-His6*: Plasmid pQE30-*eGFP* was transformed into DH5α and checked for expression, similar to *EcGrx2-eGFP-His6*, prior to microscopic studies.

### DHA reductase assay

All enzymatic reactions were analysed using a UV-Spectrometer: 3100 (Amersham / Pharmacia). DHA reductase activity of *Ec*Grx2 and its mutants *Ec*Grx2^C9S^ and *Ec*Grx2^C9S/C12S^ was assayed as described earlier^37^ with minor modifications. Prior to enzyme assay, protein aliquotes were thawed and exchanged to buffer containing 20 mM Tris-HCl pH 8.0, 150 mM NaCl to remove DTT and βME, using 3kDa Centriprep (GE). The reaction was initiated by adding 2 mM GSH into the 500 μl reaction mixture containing 0.08 μg protein in 50 mM sodium phosphate buffer pH 7.4 and 0.1 mM DHA. Absorbance of AsA was monitored at 265 nm for 5 minutes. Curve fitting and V_max_ and K_m_ were calculated using GraphPad Prism software. For inhibition assay, the reaction was initiated by adding 2 mM GSH and a fixed concentration of 45µM ZnSO_4_ (IC_50_) in 500 μl reaction mixture containing 0.08 μg of protein, with varying concentrations of DHA (0.01-9 mM). Control experiments were carried out without ZnSO_4_. Lineweaver-Burk plot was plotted using Excel (Microsoft).

### Chemical modification

*Ec*Grx2 was incubated with 0.5 to 2 mM iodoacetamide (IAM) at 4 °C for 2 h, followed by buffer exchange into 20 mM Tris-HCl pH 8.0, 150 mM NaCl using centriprep (3 kDa) to remove excess reagents.

### Thermal shift assay (TSA)

20 µM tagless *Ec*Grx2 was added to the buffer containing 20 mM Tris-HCl pH 7.5, 10 mM NaCl and 2X SYPRO orange dye (S6650, Invitrogen), and 20 µM of various metal ions (CuSO_4_, FeSO_4_, ZnSO_4_, CaCl_2_, NiSO_4_, MgSO_4_, CoCl_2_, MnCl_2_). The mixture was spun at 1000 x g prior to TSA. The samples were heated from 20 to 95 °C at the rate of 1 °C min^-1^ and fluorescence signals were monitored using quantitative real-time PCR (CFX96 Real-Time System, Bio-Rad). Each experiment was done in triplicate and the average values were taken for analysis. Assay buffer with and without added protein were used as controls.

### Crystallization and structure determination: *Ec*Grx2-GSF-Zn^2+^

Prior to setting up crystallization plates, DTT was added to a final concentration of 5 mM to the tagless protein (25 mg/ml) and incubated for 2 hr at RT. GSH and DHA were added to the above mixture to a final concentration of 20 and 10 mM, respectively, and incubated overnight at 4 °C. The protein-ligand solutions were centrifuged at ∼ 16000 x g for 15 minutes at 4 °C. Crystallization trails were done at 20 °C using sitting-drop vapor diffusion method using protein to precipitant ratio; 1:1, 1:2 and 2:1 to a final volume of 150 nl, using commercial crystallization screens (Hampton Research and Molecular Dimensions) and Mosquito robot (TTP Labtech). Plate-like crystals were obtained after 5 days in the condition containing 10 mM 0.01 M ZnSO_4_.7H_2_O, 0.1M MES monohydrate pH 6.5, 25% PEGMME 550. Crystals were quickly washed and soaked in the reservoir condition containing 20% (v/v) glycerol as a cryoprotectant and flash-cooled in LN2. Diffraction data were collected at ID29 beamline, ESRF, Grenoble, France and integrated and scaled using the autoPROC^38^. The crystal belongs to the space group P2_1_2_1_2 with one *Ec*Grx2-Zn-GSF complex per asymmetric unit. Initial phases were obtained using Molecular Replacement method, using AutoMR module in PHENIX suite^39^ with *Ec*Grx2-GSH (PDB: 4KX4) as the search model. Iterative model building and refinement were carried out in Coot^40^ and Refmac^41^. Coordinates and restraints for GSF were generated using through JLigand (CCP4i suite). The final model was refined to R_work_/R_free_ = 0.13/0.18 (Table 1). Tetrahedral coordination of Zn^2+^ was validated using CheckMyMetal server^19^ (Supplementary Table 2). ***Ec*Grx2-GSH:** Protein at 25 mg/ml was treated in a similar way as in the case of *Ec*Grx2-Zn-GSF complex, before setting up plates. GSH bound *Ec*Grx2 crystals were obtained after 5-10 days in 0.2 M ammonium acetate, 0.1 M sodium citrate tribasic dihydrate pH 5.6, and 30% PEG 4000, flash-cooled in LN2 with 20% (v/v) glycerol as cryoprotectant. Data were collected at ID30-A1 beamline (ESRF) and integrated and scaled as mentioned before. Crystals diffracted to 1.45Å and belonged to P 2_1_ space group with four molecules of *Ec*Grx2-GSH complex per asymmetric unit. Data processing, molecular replacement, model building and refinement were performed as above. The final model was refined to Rwork/Rfree = 0.138/0.160. ***Ec*Grx2-apo:** Crystals belonging to the space group P 2_1_ 2_1_ 2_1_ were grown at 35 mg/ml in 0.1 M HEPES pH 7.5, 20% w/v polyethylene glycol 10,000 and diffracted beyond 2.38 Å and X-ray data was collected at XRD2 beamline, Elettra, Trieste, Italy. Data processing and scaling were done using iMosflm^42^. Molecular replacement, model building, refinement was done as above. The final model was refined to Rwork/Rfree =0.195/0.249. Structure-based sequence alignment was made using Dali server. Figures were made using PyMol (Schrodinger).

### Molecular dynamics stimulation

Atomistic **s**imulations were performed using the *Ec*Grx2-apo and *Ec*Grx2-Zn^2+^-GSF complex structures using Schrodinger (licenced to ICGEB). First, individual structures were incorporated into Maestro. The protein preparation wizard was used to assign bond orders, addition of hydrogens, filling missing side chains, creating zero bond order to metals and creating disulphide bonds. Hydrogen bond network was optimized at pH 7.4 and crystallographic water molecules were retained during protein preparation. A final restrained minimization was performed using the OPLS3e force field. The system was built using Desmond. The orthorhombic box was solvated with TIP3P solvent model and further neutralized with 0.15 M NaCl as counter ions. Simulations were performed at 300 K and at1.0325 bar pressure using NPT ensemble. Solvated system was relaxed with a series of energy minimization before production MD. The total simulation time for each system was set to 1000 ns and the coordinates were saved at an interval of 50 ps.

### Ascorbate supplementation assay

The assay was performed using BW25113 and JW1051-3 (*grxB*^*-*^, Kan^R^) strains using a plating protocol^43^ with modifications. Briefly, cells were grown in LB broth till OD_600_ of 0.3. Aliquots consisting of 0.5 ml culture and 4.5 ml of soft agar, containing 0.8% agar in M9 media with respective antibiotics, were mixed and poured immediately over hard agar (1.5% agar in M9 minimal media, with respective antibiotics). After the plates were solidified, a 0.8 cm diameter filter paper disc (Whatman 3mm Chr), saturated with 10 μl of 6.6 M H_2_O_2_, was placed in middle of the plate and incubated at 37 °C for 12 h. Each experiment was done in triplicates with biological replicates (n=2). The zone of clearance (diameter) was calculated by taking the mean value of measurements taken from three different directions. Statistical analysis, one-way ANOVA followed by Tukey’s test, was done using SPSS (V. 20). A p-value of ≤ 0.05 is considered significant for the test.

### Macrophage killing assay

RAW 264.7 macrophages were grown in Dulbecco’s modified Eagle’s media (DMEM, cell clone) supplemented with 10% Fetal Bovine Serum (FBS, GIBCO) and maintained at 37 °C in a humified, 5% CO_2_ atmosphere. LPS (20 ng/ml) treatment was given to RAW 264.7 cells for 24 h (for pre-activation of macrophages), washed with plain DMEM and infected with respective bacteria (BW25113 and JW1051-3) with a MOI of 10, for 30 min. Thereafter, the cells were washed with 1X PBS and maintained in complete DMEM with amikacin (200µg/ml) for 1 h to kill extracellular bacteria. Subsequently, cells were washed twice with 1X PBS and maintained in complete DMEM for monitoring at different experimental time points (0, 30, 60, 120 and 240 min). At every timepoint, cells were lysed using 1X PBS containing 0.06% SDS, 1/100 dilution of the lysate was spread on plain LB plates, and incubated at 37 °C for 12 h. Plates were collected after 12 h and colony count was taken for calculating the CFU/ml.

### Confocal and TIRF-SIM microscopy

BW25113 WT cells transformed with pQE30-*grxB-eGFP* and pQE30-eGFP, expressing His6-*Ec*Grx2-eGFP and His6-eGFP, were grown at 37 °C till stationary phase, 12 h. Cells were pelleted down at 2095 x g at RT and washed twice with 1X PBS, followed by M9 minimal media to remove the autofluorescence from LB media. Cells were then incubated for 10 minutes at RT with FM4-64 plasma membrane stain (#T13320, Invitrogen) and washed using M9 minimal media. Cells were layered onto slides coated with poly-L-lysine, mounted using Fluroshield (# F6057, Sigma) and sealed. Images were acquired using a Nikon A1R-A1 fluorescence microscope with a ×100 objective with 4.1 X zoom factor. The same field of view was chosen for TIRF-SIM imaging. Membrane localization of *Ec*Grx2 in BW25113 was observed using 100X objective and a SIM based illumination using ±1 order light with a fixed critical angle for TIRF Imaging. The light source was a 488 nm laser. Fluorescence was observed through a 488 nm band-pass filter using an electron-multiplying CCD (iXon 897, Andor). The beam was totally internally reflected at an angle of 63° from the surface normal. An elliptical area of 32× 32 µm was illuminated for TIRF-SIM imaging. The images were processed using NIS-Elements AR software.

### Membrane isolation and detergent extraction

BL21 (DE3) cells transformed with pET16b-*grxB-eGFP*, expressing *Ec*Grx2-eGFP-His6, were resuspended at 1:10 ratio (pellet:buffer) in lysis buffer containing 50 mM Tris-HCl pH 8, 150 mM KCl, 10 Mm MgCl_2,_ 0.2 mgml^-1^ Lysozyme, 3 unitsµl^-1^ DNase-I, 1 mM Benzamidine, 1 mM PMSF were disrupted by sonication. Cell lysate was centrifuged at 15000 g for 20 min at 4 °C. The supernatant was spun at 1,00,000 x g for 1 h at 4 °C. The pellet containing pure membrane was resuspended at 1:10 ratio in wash buffer containing 50 mM Tris-HCl pH 8.0, 5 % Glycerol, 100 mM KCl, 1 mM PMSF and 2 M Urea, using a Dounce homogenizer, and left at 4 °C for overnight. Sample was spun again at 1,00,000 g at 4 °C for 1 h. The pellet was further washed with resuspension buffer containing 50 mM Tris-HCl pH 8.0, 5 % Glycerol, 100 mM KCl and 1 mM PMSF, at 1:10 ratio, and spun at 1,00,000 x g at 4 °C for 1 h. Equal amount of washed membrane pellet was solubilized in the above buffer containing 1% DM, DDM, OG, LDAO and SDS, respectively, and incubated at RT for 1 h and spun at 1,00,000 x g for 1 h. Supernatant containing detergent extracted *Ec*Grx2 were subjected to reduced and denaturing SDS-PAGE and western blot analysis using anti-His antibodies (A7058, sigma). Equal volume of ultra-supernatant was loaded on the gel. SDS: sodium dodecyl sulphate, OG: n-Octyl-β-D-glucopyranoside, DDM: n-dodecyl-β-D-maltopyranoside, DM: n-Decyl-β-D-Maltopyranoside, LDAO: Lauryl dimethylamine oxide.

### Bilayer electrophysiology

Planar lipid bilayer bilayers were formed using 1,2-diphytanoyl-*sn*-glycero-3-phosphocholine (DPhPC, Avanti Polar Lipids) lipids by employing Montal and Muller technique. Lipid bilayers were formed across ∼100 µm in diameter an aperture in a 25-µm thick polytetrafluoroethylene (Teflon) film (Goodfellow, Cambridge). Notably, Teflon film separated the bilayer cuvette made in Delrin into cis and trans chambers (500 µL each). Bilayers were formed by pre-painting on each side of the Teflon aperture with hexadecane in n-pentane (1 μL, 5 mg mL^-1^). Then, both cis and trans chambers were filled with 1 M KCl, 10 mM MES, pH 6.0 that acts as the electrolyte. Finally, DPhPC lipids in n-pentane (2 μL, 5 mg mL^-1^) were added to both sides of the chamber and a solvent-free lipid bilayer was formed by lowering and raising the electrolyte buffer subsequently bringing the two lipid surface monolayers at the aperture. The purified *Ec*Grx2 was added to the cis side of the bilayer chamber with 1 mM DTT and a voltage of + 100 mV was applied to facilitate the channel insertion. The cis chamber was connected to the grounded electrode and the trans chamber was attached to the working live electrode. A potential difference was applied through a pair of Ag/AgCl electrodes, set in 3% agarose containing 3.0 M KCl. The current was amplified by using an Axopatch 200B amplifier, digitized with a Digidata 1550B and recorded with the pClamp 10.6 acquisition software (Molecular Devices, CA) with a low-pass filter frequency of 2 kHz and a sampling frequency of 10 kHz. The data were analysed and prepared for presentation with pClamp (version 10.6, Molecular Devices, CA) and Origin 9.0.

### Phylogenetic tree construction

HMMER 3.3.3 was used to retrieve *Ec*Grx2 like sequences with *E. coli* str. K-12substr. MG1655 Grx2 as input from UniProtKB database. *grxB* gene encoded Grx2 sequences were manually selected (one per genus) with the highest E-value and having two domain architecture similar to *Ec*Grx2. A total of 152 sequences was used for multiple sequence alignment using MUSCLE. The aligned sequences were then given as input to MEGA-X^44^ to construct a phylogenetic tree using maximum likelihood method with default settings. The tree with the highest log likelihood (−40838.84) was used to make the circular phylogenetic tree using iTOL^45^.

## Supporting information

Fasta sequences used for phylogenetic tree construction

## Acknowledgments

Authors thank H. Jane Dyson, The Scripps Research Institute, La Jolla, for *Ec*Grx2 construct; MK Reddy for valuable inputs, Purnima Kumar for help with confocal microscopy; Mahavir Tanwar for TIRF-SIM imaging; Aravind Penmatsa for pET16b-eGFP vector; Bichitra Kumar Biswal and Ravikant Pal for preliminary X-ray data collection at National Institute of Immunology; Lakshminarayan M Iyer, NCBI, NIH for inputs with phylogenetic analysis and ciritical reading of the manuscript; Raghurama P. Hegde and Annie Heroux for beamline support at Elettra (Access to the XRD2 beamline at the Elettra synchrotron, Trieste was made possible through grant-in-aid from the Department of Science and Technology, India, vide grant number DSTO-1668); Data collection at ESRF beamlines ID30A-3 and ID29 was facilitated by the ESRF Access Program of the Regional Centre for Biotechnology, supported by the Department of Biotechnology, Government of India vide grant number BT/PR36150/INF/22/214/2020;

## Funding

SNS was supported by DST-INSPIRE fellowship (IF150373) Department of Biotechnology (BT/PR8766/BRB/10/1701/2018) Department of Biotechnology (BT/PR28080/BID/7/836/2018) Department of Science and Technology (EMR/2017005066) ICGEB Core funds.

## Author contributions

SNS, BKD and RR generated constructs, expressed, purified *Ec*Grx2, performed enzyme assays. PR performed initial expression and purification. SNS performed ascorbate supplementation and SK and AC performed statistical analysis. SNS, AK, RR and AA solved the crystal structures and SS performed MD simulation. SU and SNS performed macrophage killing assay, supervised by DK. NP and SNS carried out ion-channel recordings, supervised by KRM. P. Ray and AC provided inputs to experimental plan and data analysis. SNS and AA wrote the manuscript with inputs from all the authors. AA conceived, planned and orchestrated the project.

## Competing interests

Authors declare that they have no competing interests.

## Data availability

The atomic coordinates and structure factors are available from Protein Data Bank under the accession codes 7DKR, 7DKP, 7D9L for native, GHS-bound and Zn^2+^ inhibited structures, respectively. The raw X-ray diffraction data is available from Integrated Resource for Reproducibility in Macromolecular Crystallography (IRRMC) repository: https://proteindiffraction.org/

**Supplementary Figure 1.**
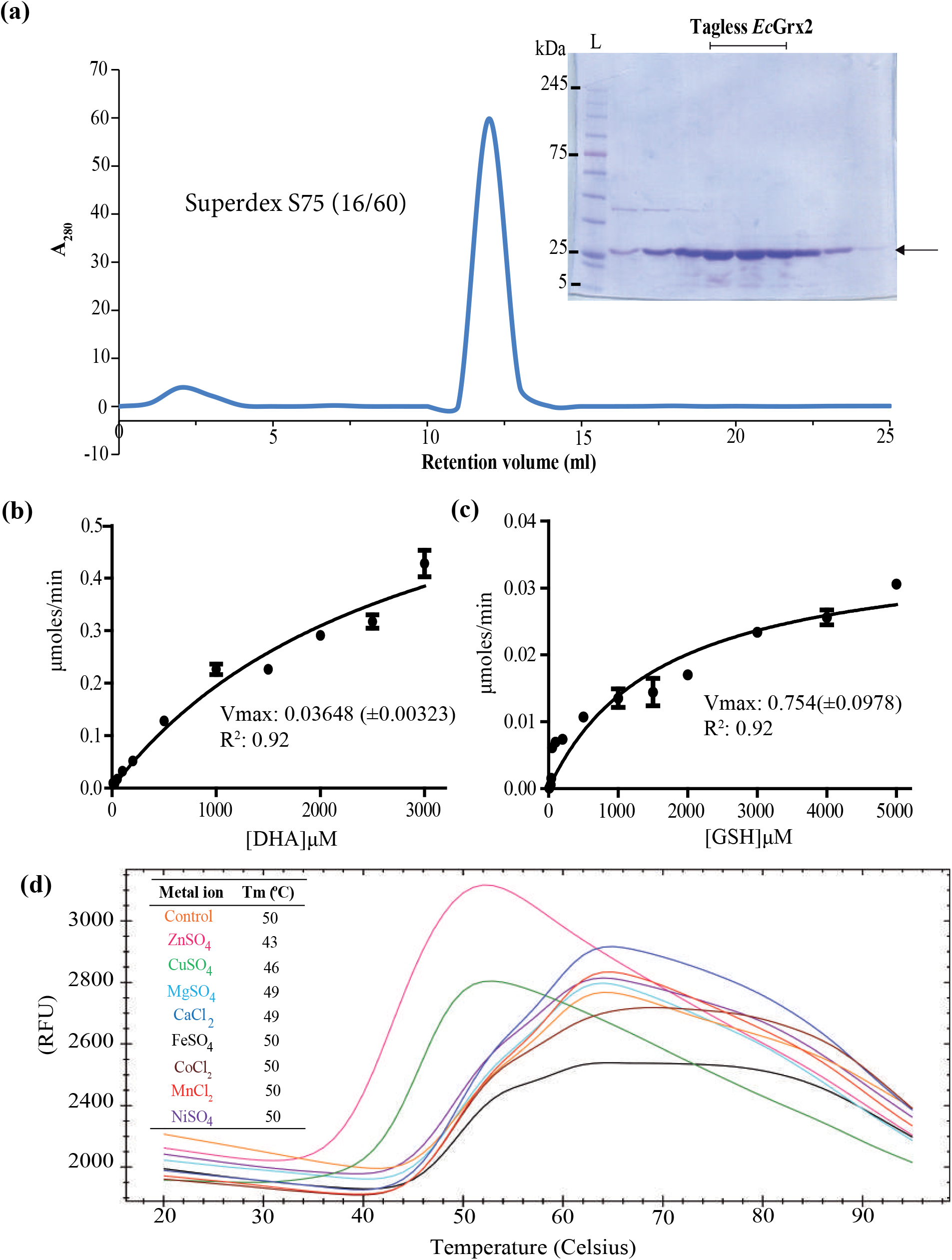
Purification, DHA reductase activity and thermal stability assays with *Ec*Grx2. **(a)** Chromatogram from Superdex S-75 (16/60) column with inset showing peak fractions loaded onto 4-20 % SDS-PAGE showing purified *Ec*Grx2 at ∼24kDa size. **(b)** Michaelis-Menten graphs of *Ec*Grx2 showing DHA reductase activity. AsA min^-1^ monitored at A_265_ plotted against different concentration of DHA (μM) and **(c)** GSH (μM). **(d)** Identification of divalent metal ion binding to *Ec*Grx2 by thermal shift assay. Inset table shows comparison of melting temperatures (T_m_) from thermal shift assay. Data shown represent three technical replicates. Different divalent metal ions are indicated with different colours.

**Supplementary Figure 2.**
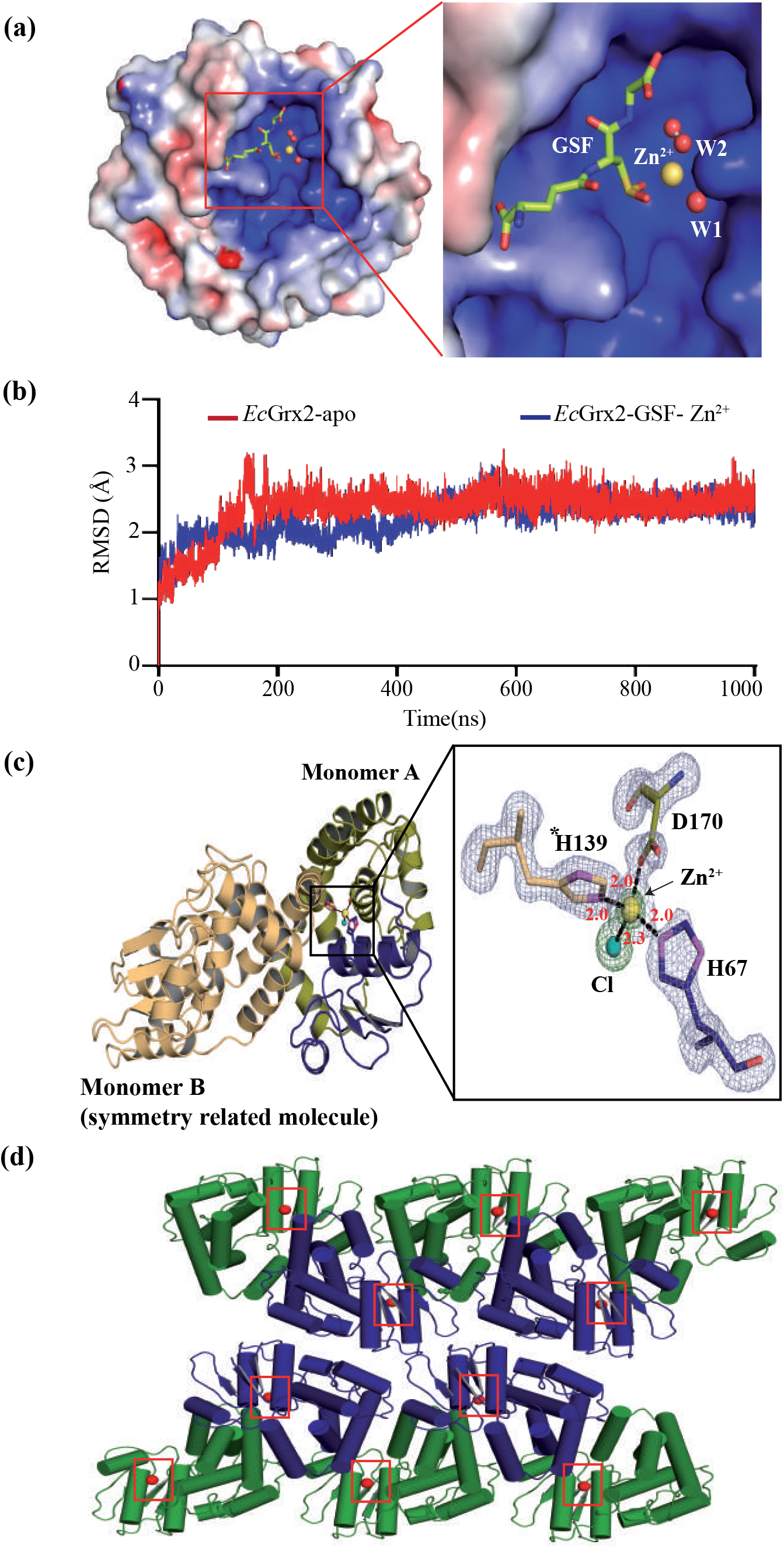
Structural analysis of Zn^2+^ inhibited complex crystal structure. **(a)** Electrostatic surface map of the active site showing Zn^2+^ inhibited state. Positively charged region is shown in blue while negatively charged region is in red. **(b)** Backbone RMSD profile of *Ec*Grx2-apo and *Ec*Grx2-GSF-Zn^2+^ complex during 1 µs MD simulation is shown. **(c)** Second Zn^2+^ binding site formed at the interface of symmetry related monomers: A and B with inset showing the tetrahedral coordination formed by A-His67 (blue), A-Asp170 (pea), Chloride ion (teal) and B-His139 (sand). The omit difference (Fo-Fc) map is contoured at 3σ with Zn^2+^ and Cl^-^ ions shown at 15σ and 3σ, respectively. **(d)** Crystal packing showing the second Zn^2+^ ion.

**Supplementary Figure 3.**
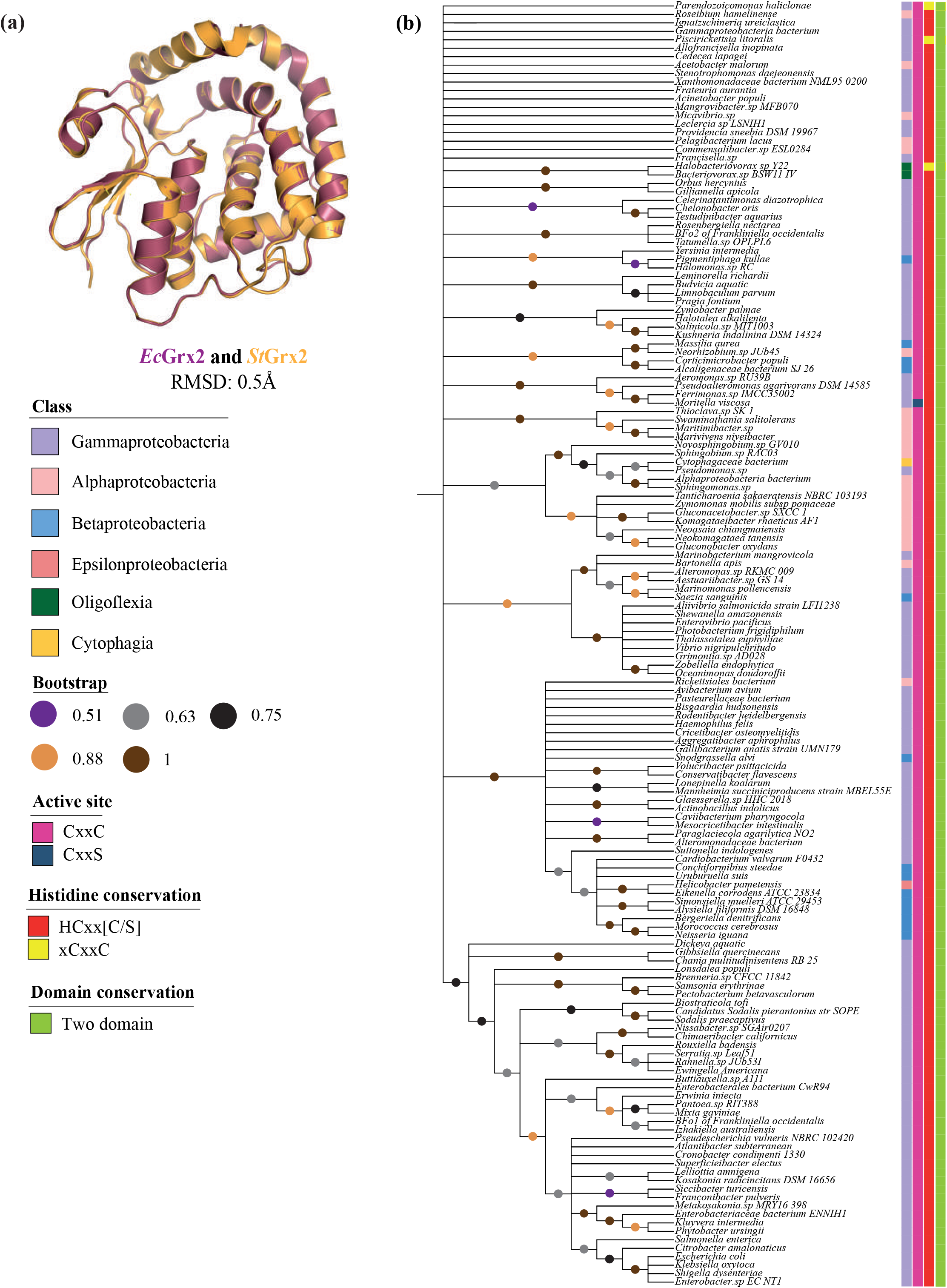
Structural homology and distribution of *grxB* gene in Gram-negative bacterial genomes. **(a)** Structural superposition of *Ec*Grx2 (7DKP, violet) with *St*Grx2 (3IR4, orange) showing very low RMSD. **(b)** Phylogenetic tree of Gram-negative bacteria possessing *grxB* encoded two domain Grx2 is given. Bootstrap value=1000.

**Supplementary Table 1.** Dehydroascorbate reductase activity data of *Ec*Grx2 and its mutants.

|  | *EcGrx2* * | *EcGrx2*<br>**C9SC12S\*** |
| --- | --- | --- |
| K<sub>m</sub>DHA (μM) | 2879<br>(±640.4) | 968<br>(±462.3) |
| K<sub>m</sub>GSH (μM) | 1630<br>(±363) | ND |
| Specific activity<br>(μM<sup>-1</sup>min<sup>-1</sup>mg) | 1.8845<br>(±.01) | 0.09705 |
| K<sub>cat</sub>/K<sub>m</sub> (DHA)<br>(M<sup>-1</sup>min<sup>-1</sup>) | 7.9704 | 1.22090 |
| Reference | This study | This study |

\*Mean± standard deviation of three separate set of experiments

**Supplementary Table 2.** CheckMyMetal (CMM)-report for *Ec*Grx2-GSF-Zn^2+^ complex.

| Warning: Coordinating ligands by symmetry operation are labeled with prefix 'sym-' |  |  |  |  |  |  |  |  |  |  |  |  |
| --- | --- | --- | --- | --- | --- | --- | --- | --- | --- | --- | --- | --- |
| ID | Res. | Metal | Occupancy | B factor (env.)<sup>1</sup> | Ligands | Valence | nVECSUM | Geometry | gRMSD (°)<sup>1</sup> | Vacancy<sup>1</sup> | Bidentate | Alt. metal |
| A:216 | ZN | ZN | 1 | 13.1 (15.1) | O<sub>4</sub>N<sub>1</sub> | <u>1.6</u> | <u>0.18</u> | Tetrahedral | 6.3° | 0 | 1 | Cu, Co |
| A:217 | ZN | ZN | 1 | 19 (21) | O<sub>1</sub>N<sub>2</sub> | 1.9 | 0.045 | Tetrahedral | 3.8° | <u>25%</u> | 0 |  |
| A:218 | CL | CL | 1 | <u>26.8</u> (22.6) |  | N/A | N/A | Free | N/A | N/A | N/A |  |
| Legend: |  |  | Not applicable | Outlier | Boderline | Acceptable |  |  |  |  |  |  |

| Column | Description |
| --- | --- |
| Occupancy | Occupancy of ion under consideration |
| B factor (env.)<sup>1</sup> | Metal ion B factor, with valence-weighted environmental average B factor in parenthesis |
| Ligands | Elemental composition of the coordination sphere |
| Valence<sup>2</sup> | Summation of bond valence values for an ion binding site. Valence accounts for metal-ligand distances |
| nVECSUM<sup>3</sup> | Summation of ligand vectors, weighted by bond valence values and normalized by overall valence. Increase when the coordination sphere is not symmetrical due to incompleteness. |
| Geometry<sup>1,4</sup> | Arrangement of ligands around the ion, as defined by the NEIGHBORHOOD algorithm |
| gRMSD(°)<sup>1</sup> | R.M.S. Deviation of observed geometry angles (L-M-L angles) compared to ideal geometry, in degrees |
| Vacancy | Percentage of unoccupied sites in the coordination sphere for the given geometry |
| Bidentate | Number of residues that form a bidentate interaction instead of being considered as multiple ligands |
| Alt. metal | A list of alternative metal(s) is proposed in descending order of confidency, assuming metal environment is accurately determined. This feature is still experimental. It requires user discrimination and cannot be blindly accepted |

**Supplementary Table 3.** Statistical analysis of organisms in ascorbate supplementation assay.

| **TUKEY HSD (one way ANOVA)** |  |  |  |  |
| --- | --- | --- | --- | --- |
| **Dependent variable** | **(I) Groups** | **Dependent variable** | **(J) Groups** | **Sig.** |
| JW1051-3 | 0% ascorbate | JW1051-3 | 0.3% ascorbate | 0.000 |
| BW25113 | 0% ascorbate | BW25113 | 0.3% ascorbate | 0.000 |
| BW25113 | 0% ascorbate | JW1051-3 | 0.3% ascorbate | 0.000 |

\*The mean difference is significant at 0.05 level.

**Supplementary Table 4.** Statistical analysis of bacterial survival in LPS activated RAW264.7.

| **TUKEY HSD (one way ANOVA)** |  |  |  |
| --- | --- | --- | --- |
| **Dependent variable** | **(I) Groups** | **(J) Groups** | **Sig.** |
| 0 min | JW1051-3 | BW25113 | 0.92 |
|  | BW25113 | JW1051-3 | 0.92 |
| 30 min | JW1051-3 | BW25113 | 0.98 |
|  | BW25113 | JW1051-3 | 0.98 |
| 60 min | JW1051-3 | BW25113 | 0.18 |
|  | BW25113 | JW1051-3 | 0.18 |
| 120 min | JW1051-3 | BW25113 | 0.1 |
|  | BW25113 | JW1051-3 | 0.1 |
| 240 min | JW1051-3 | BW25113 | <0.001 |
|  | BW25113 | JW1051-3 | 0 |

\*The mean difference is significant at 0.05 level.

**Supplementary Table 5.**
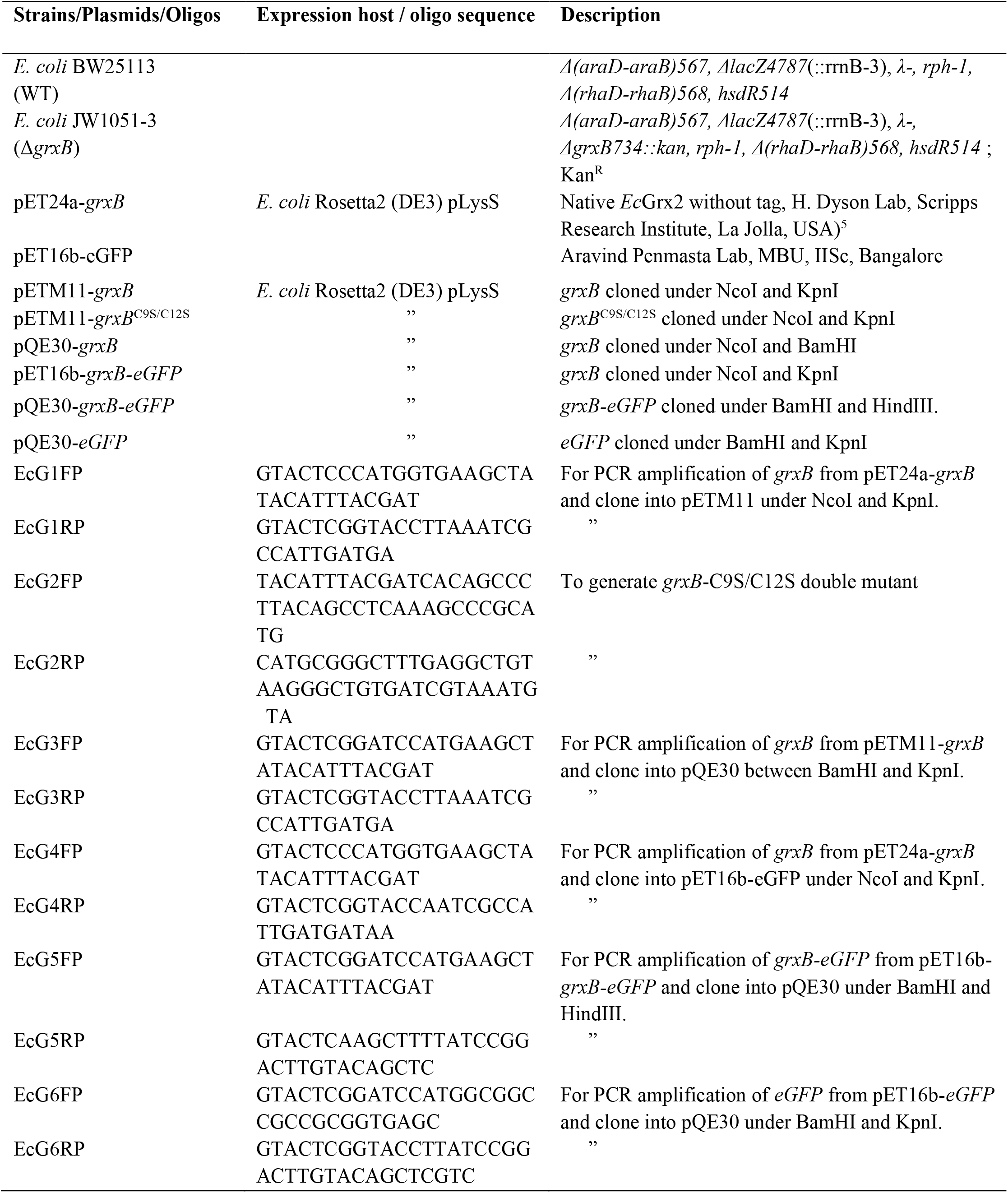
Plasmids, *E. coli* strains, and oligos used in this study.

**Supplementary Table 6.** Composition of buffers used for protein purification and membrane extraction.

| **Name of the buffer** | **Composition** | **pH** | **NaCl (mM)** | **KCl (mM)** | **Imidazole (mM)** | **βME (mM)** | **DTT (mM)** | **Glycerol (%)** | **PMSF (mM)** |
| --- | --- | --- | --- | --- | --- | --- | --- | --- | --- |
| A* | 50 mM Tris-HCl | 7.5 | 10 |  |  | 3 |  |  | 1 |
| B | 10 mM MES | 6.5 | " |  |  |  | 3 |  |  |
| C | 20 mM Tris-HCl | 8.0 | 1000 |  |  | 3 |  |  |  |
| D | " | " | 150 |  |  | 3 |  |  |  |
| E | 25 mM Tris-HCl | " | " |  | 10 |  |  |  |  |
| F | " | " | " |  | 10 |  |  |  |  |
| G | " | " | " |  |  |  |  |  |  |
| H | " | " | " |  | 300 |  |  |  |  |
| I | 20 mM Tris-HCl | " | " |  |  |  |  |  |  |
| J# | 50 mM Tris-HCl | " |  | 150 |  |  |  |  | 1 |
| K\$ | " | " | | 100 | | | | 5 | 1 |
| L | " | " |  | 100 |  |  |  | 5 | 1 |

\*Contains 1 mM Benzamidine-HCl, 3 unitsμl<sup>-1</sup> DNase I and 0.2 mg ml<sup>-1</sup> Lysozyme, #-contains 10mM MgCl<sub>2</sub>, 1 mM Benzamidine-HCl, 3 unitsμl<sup>-1</sup> DNase I and 0.2 mg ml<sup>-1</sup> Lysozyme, \$- contains 2M Urea.

**Supplementary Video 1.**
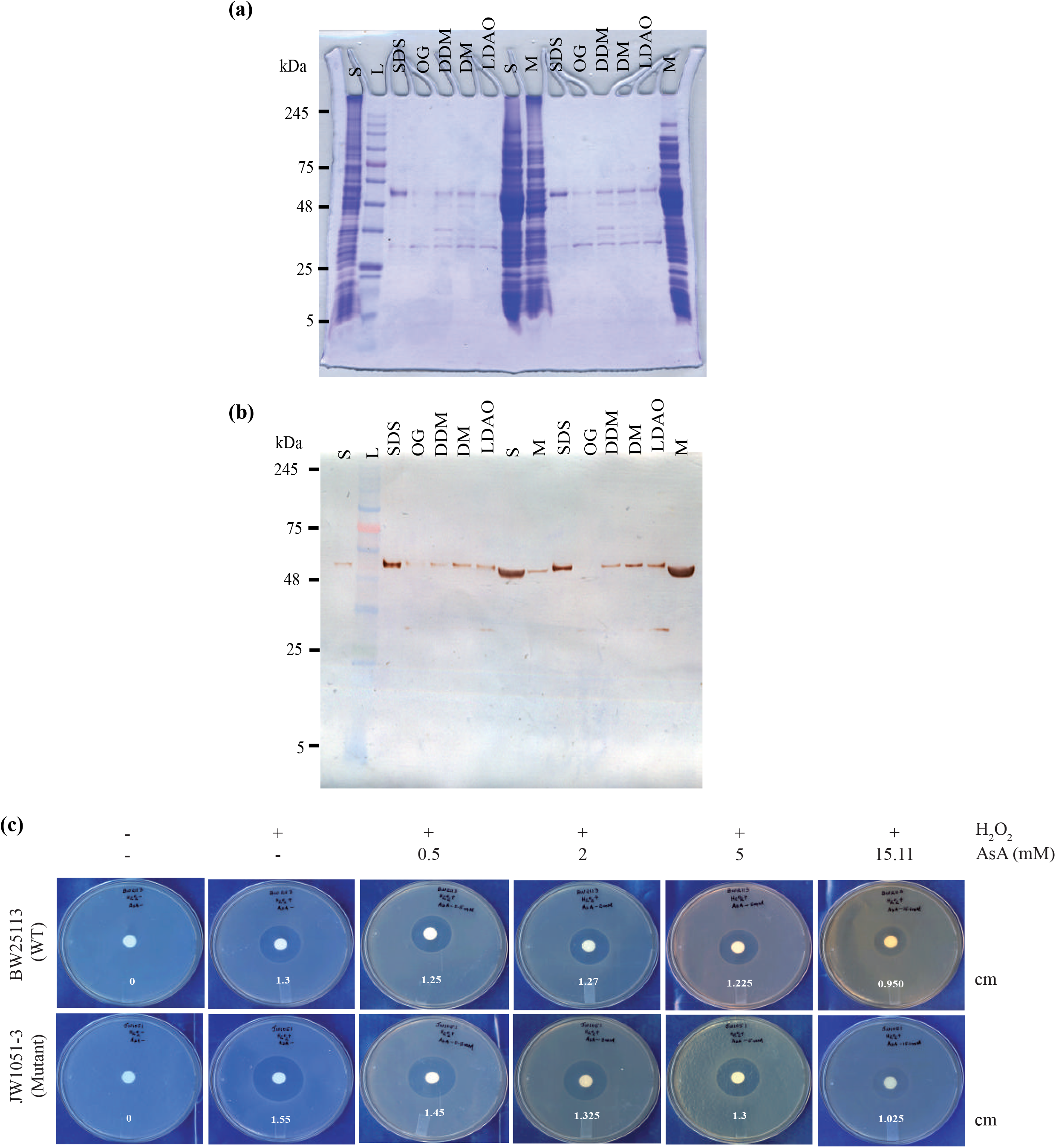
MD simulation trajectory showing Zn^2+^, coordinated waters (W1, W2), GSF, and His8 of *Ec*Grx2. **(a)** Uncropped SDS-PAGE after membrane isolation and detergent extraction, **(b)** Corresponding western blot of (a), and **(c)** Scanned full images of petri plates used for making Figure 2a. Strains BW25113 and JW1051-3 are labelled, by hand, as BW2113 and JW1051, respectively.

## Notes

### Competing Interest Statement

The authors have declared no competing interest.

